# TOL proteins mediate vacuolar sorting of the borate transporter BOR1 in *Arabidopsis thaliana*

**DOI:** 10.1101/342345

**Authors:** Akira Yoshinari, Barbara Korbei, Junpei Takano

## Abstract

Boron (B) is an essential micronutrient for plants, however, it shows cytotoxicity at high concentrations. A borate transporter BOR1 is required for efficient transport of boron (B) toward the root stele in *Arabidopsis thaliana*. BOR1 shows polar localization in the plasma membrane of various root cells toward the stele-side under B limitation. To avoid over-accumulation of B, BOR1 in the plasma membrane is rapidly internalized and transported into the vacuole for proteolysis after high-B supply in an ubiquitination-dependent manner. Although BOR1 has been predicted to be transported into multi-vesicular bodies/late endosomes (MVB/LEs) via the endosomal sorting complex required for transport (ESCRT) machinery, experimental evidence was absent so far. In this study, we investigated the intracellular localization of BOR1 by visualizing endomembrane compartments, and tested the involvement of ESCRT-0-like proteins TOM1-LIKEs (TOLs) in the vacuolar sorting of BOR1. Under low-B conditions, a large portion of cytoplasmic BOR1 was localized in the *trans*-Golgi networks/early endosomes (TGN/EEs) labeled with VHA-a1 subunit. Pharmacological interference of endosomal recycling using brefeldin A induced colocalization of BOR1 with RabA5D, which labels recycling vesicles associated with the TGN. These data suggest that BOR1 cycles between plasma membrane and TGN/EE via RabA5D-positive endomembrane compartments under low-B conditions. On the other hand, under high-B conditions, BOR1 was localized in the inside of TOL5-positive MVB/LEs. To examine the roles of TOL proteins in intracellular trafficking of BOR1, we analyzed BOR1-GFP localization in the TOL quintuple mutant (*tolQ*; *tol2-1tol3-1tol5-1tol6-1tol9-1*) after high-B supply. In the *tolQ* mutant, vacuolar sorting of BOR1 was delayed, while the polar localization of BOR1 was not disturbed. Taken together, BOR1 is constantly transported to the TGN/EE by endocytosis and recycled to the plasma membrane likely via RabA5D-positive endomembrane compartments under low-B conditions. On the other hand, BOR1 is transported to the vacuole via TOL5-positive MVB/LEs under high-B conditions. TOL proteins are required for sorting of ubiquitinated BOR1 into MVB/LE for vacuolar degradation but not for the polar trafficking of BOR1.

## 1. INTRODUCTION

Plants take up mineral nutrients in soil solution via transporters localized in the plasma membrane of root cells. Most nutrients, however, exert toxic effects to plant cells when excessively accumulated. Due to the sessile lifestyle, higher plants regulate abundance and localization of a variety of transporters in response to the nutrient availability in the local environment. Subcellular localization and degradation of plasma membrane-localized transporters are regulated through the membrane trafficking system. Generally, plasma membrane proteins are translated in the rough endoplasmic reticulum (ER), and transported over the Golgi apparatus and *trans*-Golgi network/early endosome (TGN/EE) to the plasma membrane. The endocytosed proteins from the plasma membrane are targeted to the TGN/EE and subsets are recycled to the plasma membrane from the TGN/EE. On the other hand, plasma membrane proteins with ubiquitination are sequestered into intralumenal vesicles (ILVs) of multi-vesicular body/late endosome (MVB/LE). Subsequently, MVB/LEs fuse with lytic vacuoles and ILVs are released into the vacuole for proteolysis (Paez Valencia et al. 2016).

Boron (B) is an essential micronutrient for plants, but it is toxic when accumulated in excess (Shorrocks, 1997; Nable et al. 1997; Sakamoto et al. 2013). Under low-B conditions, the boric acid channel NIP5;1 and the borate exporter BOR1 show polar localization in the plasma membrane of root cells toward the soil and stele side, respectively (Takano et al. 2002, 2006, 2010). The polar localization of NIP5;1 was shown to be important for efficient B-transport from the root surface to shoots (Wang et al. 2017). Although it has not been clearly demonstrated, the polar localization of BOR1 has been considered to be important for the efficient translocation of B (Yoshinari and Takano, 2017). The polar localization of BOR1 is maintained by intracellular trafficking, and it requires YxxΦ (Y, tyrosine; x, any residues; Φ, bulky hydrophobic residues) motifs and a D/ExxxLL/I (D/E, aspartate or glutamate; x, any residues; L, leucine; L/I, leucine or isoleucine) motif in the cytoplasmic region of BOR1 (Takano et al. 2010; Yoshinari et al. 2012; Wakuta et al. 2015). BOR1-GFP protein accumulated in intracellular aggregates upon treatment with brefeldin A (BFA), an inhibitor of endosomal recycling. indicating that BOR1 undergoes constitutive endocytosis. In Arabidopsis, DYNAMIN-RELATED PROTEIN 1A (DRP1A), which is involved in clathrin-mediated endocytosis, is required for constitutive endocytosis and polar localization of BOR1 (Yoshinari et al. 2016).

Arabidopsis RabA and RabE GTPases are homologs of Rab8 and Rab11 in mammals, respectively (Rutherford and Moore, 2002). In mammalian cells, Rab8 and Rab11 GTPases coordinately play roles in exocytic vesicular trafficking (Knödler et al. 2010; Mizuno-Yamasaki et al. 2012). Consistently, in *Nicotiana benthamiana*, dominant-negative inhibition of RabA1B inhibited the secretory trafficking of BOR1 to the plasma membrane (Choi et al. 2013). Additionally, pharmacological interference of RabA and RabE GTPases by Endosidin 16 dramatically inhibited the secretory trafficking of BOR1 (Li et al. 2017). These results suggested that RabA and RabE GTPases are involved in exocytic trafficking events including the secretion and the endosomal recycling in plant cells.

Under high-B conditions, BOR1 is down-regulated by endocytic degradation and translational repression to avoid over-accumulation of B in shoots (Takano et al. 2005; Aibara et al. 2018). BOR1 accumulated in the plasma membrane is first rapidly transferred into the multi-vesicular bodies/late endosomes (MVB/LEs) labeled with a Rab5 GTPase homolog, ARA7, and then to the vacuoles upon high-B supply (Takano et al. 2005; Viotti et al. 2010). Intriguingly, ADP-ribosylation factor guanine nucleotide exchange factor (ARF-GEF) BIG3 and its homologs are required for high B-induced degradation of BOR1, suggesting that ARF-GEF-dependent vesicle trafficking is involved in the vacuolar transport of BOR1 (Richter et al. 2014).

Ubiquitination has been shown to play roles in the proteolysis of plant membrane proteins including the auxin efflux transporter PIN-FORMED 2 (PIN2), the flagellin receptor FLAGELLIN-SENSING 2 (FLS2), the chitin receptors CHITIN ELICITOR RECEPTOR KINASE 1 (CERK1) and LYSIN MOTIF RECEPTOR KINASE 5 (LYK5), the iron influx transporter IRON-REGULATED TRANSPORTER 1 (IRT1), the brassinosteroid receptor BRI1, and BOR1 in Arabidopsis (Abas et al. 2006; Gohre et al. 2008; Gimenez-Ibanez et al. 2009; Barberon et al. 2011; Kasai et al. 2011; Martins et al. 2015; Liao et al. 2017; Yamaguchi et al. 2017). The flagellin-induced immune signaling is attenuated through polyubiquitination of FLS2 by the U-box E3 ubiquitin ligases PUB12 and PUB13 and subsequent vacuolar sorting (Lu et al. 2011). In the case of PIN2, polyubiquitination and/or multi-monoubiquitination are required not only for constitutive protein turnover, but also for gravistimulation-induced asymmetric degradation, auxin-induced degradation, and dark-induced degradation of PIN2 (Leitner et al. 2012). Intriguingly, ubiquitination of PIN2 is also involved in its constitutive endocytosis. In the case of BRI1, polyubiquitination promotes its endocytosis and vacuolar sorting (Martins et al. 2015).

Generally, ubiquitinated membrane proteins are targeted to the vacuole via the Endosomal Sorting Complex Required for Transport (ESCRT) machinery. The ESCRT machinery is required for the recognition and transport of ubiquitinated proteins into ILVs of MVB/LE. The ESCRT machinery consists of four subcomplexes named ESCRT-0, I, II, and III in animal and yeast. Plants lack canonical ESCRT-0 components required for the initial recognition of ubiquitinated proteins (Gao et al. 2017). A recent study clarified that TOM1-LIKE (TOL) proteins function as ESCRT-0-like components in plant cells. The Arabidopsis genome has nine TOL genes (TOL1–TOL9), which play redundant roles in vacuolar sorting of PIN2 and KNOLLE syntaxin (Korbei et al. 2013). TOL5 is localized in both cytosolic and membrane fractions, while TOL6 preferentially localizes to the plasma membrane and endomembrane compartments (Korbei et al. 2013).

In the present study, we investigated the intracellular trafficking pathway of BOR1 using fluorescent endosomal markers and multiple *tol* mutants. We found that BOR1 is transported to TOL5-positive MVB/LEs under high-B conditions and multiple TOL proteins are required for vacuolar sorting of BOR1. Our findings provide insights into vacuolar sorting mechanisms of ubiquitinated nutrient transporters in plant cells.

## 2. MATERIALS AND METHODS

### 2.1. Plant materials

*Arabidopsis thaliana* Col-0 ecotype harboring the *pBOR1:BOR1-GFP* construct (Takano et al. 2010; Yoshinari et al. 2016) and organelle marker constructs *mCherry-Got1p* (Wave18R), *mCherry-RabE1D* (Wave27R), *mCherry-RabA5D* (Wave24R), *mCherry-RHA1* (Wave7R, Geldner et al 2009), *VHA-a1-mRFP* (Viotti et al. 2010), *pTOL5:TOL5-mCherry* (Korbei et al. 2013) were generated by crossing. Plants in F1 or F2 generation were used for confocal imaging. The *tol23569* (*tolQ*) was described previously (Korbei et al. 2013). The *pBOR1:BOR1-GFP* construct was introduced into the *tolQ* mutant by floral-dip transformation.

### 2.2. Growth conditions and drug treatment

Plants were grown on solid, modified MGRL medium containing 0.5 µM boric acid (Low-B medium, Yoshinari et al. 2016) for 4 days in a growth chamber at 22°C under 16/8-h light/dark conditions. For colocalization analysis between BOR1-GFP and organelle markers, whole 4-day-old seedlings were incubated in liquid low-B medium containing 50 µM cycloheximide (CHX, 25 mM stock solution, diluted in water) for 30 min. Then plants were incubated in low-B medium containing 50 µM CHX and 50 µM BFA (50 mM stock solution, diluted in DMSO). For BFA treatment with wortmannin (Wm), 4-day-old seedlings were incubated in liquid low-B medium containing 50 μM CHX for 30 min followed by 50 M CHX, 30 M Wm, and 50 μM BFA for 120 min. For Wm treatment with high-B, plants were incubated in liquid low-B medium containing 30 µM Wm for 30 min and then transferred into liquid medium containing 100 µM boric acid and 30 µM Wm for 1 h. To analyze the endocytic rate of BOR1-GFP, seedlings were stained with 2 µM FM4-64 (*N*-(3-Triethylammoniumpropyl)-4-(6-(4-(Diethylamino) Phenyl) Hexatrienyl) Pyridinium Dibromide) diluted in liquid low-B medium containing 50 µM CHX for 30 min and subsequently treated with liquid low-B medium containing 50 µM CHX and 50 µM BFA without FM4-64 for 90 min. For visualization of endosomes, seedlings were stained with 4 µM FM4-64 diluted in liquid low-B medium for 40 min before observation.

### 2.3. Microscopy

Confocal image acquisition was performed using a Leica TCS SP8 system equipped with a DMI6000B inverted microscope and two HyD hybrid detectors using x40 water immersion (NA=1.10) and x63 glycerol immersion (NA =1.30) objective lenses (Leica Microsystems). Laser excitation/spectral detection bandwidths were 488/500–530 nm for GFP, 552/580–700 nm for mCherry and mRFP, and 488/650–700 nm for FM4-64.

### 2.4. Quantitative analysis

Quantification of fluorescent images was performed with the Fiji software (Schindelin et al. 2012; http://fiji.sc/Fiji). The Person’s correlation between GFP and mRFP/mCherry signals in cytoplasm was calculated by using the PSC colocalization plugin (French et al. 2008) with a default threshold of 40 (https://www.cpib.ac.uk/tools-resources/software/psc-colocalization-plugin/). The polarity index of BOR1-GFP was calculated as described previously (Yoshinari et al. 2016).

## 3. RESULTS

### 3.1. BOR1 is localized in the plasma membrane and the *trans*-Golgi network under low-B conditions

A previous report using immune-electron microscopy demonstrated that BOR1 is distributed to the TGN/EE in addition to the plasma membrane under low-B conditions, whereas high B concentrations (high-B) induce translocation of BOR1 into the TGN and MVBs (Viotti et al. 2010). To further elucidate the localization and trafficking of BOR1 under low-B conditions, we first observed the intracellular distribution of BOR1-GFP in root epidermal cells by confocal imaging. In the cell cortex of epidermal cells, BOR1-GFP was localized in punctate structures under a low-B (0.5 µM) condition (Supplemental Figure 1). Shortly after staining with the fluorescent endocytic dye FM4-64, BOR1-GFP and FM4-64 signals showed partial overlap in the cytoplasm. It has been shown that short-term staining with FM4-64 highlights the TGN/EE, the first destination of endocytosed lipids (Dettmer et al. 2006). Therefore, this result suggests that BOR1 exists in a subset of the TGN/EE and also other endomembrane compartments.

To further characterize the endomembrane compartments where BOR1-GFP was localized, we analyzed the colocalization between BOR1-GFP and fluorescent markers for the Golgi apparatus (Got1p homolog), TGN/EE [vacuolar H^+^-ATPase a1 subunit (VHA-a1)], secretory/recycling-associated endomembrane compartments (RabE1D and RabA5D) and MVB/LE (RHA1/RabF2A) (Sohn et al. 2003; Dettmer et al. 2006; Geldner et al. 2009; Choi et al. 2013; Li et al. 2017; Figure 1). BFA inhibits the ARF-GEF function and induces aggregation of endomembrane compartments. Because the responses of distinct endomembrane compartments to BFA are different, BFA treatment enables us to distinguish endomembrane compartments (Geldner et al. 2009). To dissect early secretory and recycling pathways, plants were co-treated with cycloheximide (CHX), an inhibitor of de novo protein synthesis, with BFA. The solvent DMSO was used as control treatment. BOR1-GFP-positive punctate structures showed close proximity to the Golgi apparatus labeled with a mCherry-Got1p homolog under control conditions (Figure 1A). After the BFA treatment, BOR1-GFP was accumulated in the core of BFA compartments, whereas the Golgi apparatus labeled with a mCherry-Got1p homolog was preferentially distributed in the periphery of the BFA compartments (Figure 1B). In the control condition, BOR1-GFP and VHA-a1-mRFP, a TGN/EE marker showed colocalization in punctate structures (Figure 1C). This observation is consistent with a previous report (Viotti et al. 2010) and the colocalization between BOR1-GFP and FM4-64 (Supplemental Figure 1). After the BFA treatment, BOR1-GFP and VHA-a1-mRFP were accumulated in the core of the BFA compartments (Figure 1D). In addition, VHA-a1-mRFP showed punctate structures around the core of BFA compartments (Figure 1D, arrows). It was in contrast to a relatively homogenous accumulation of BOR1-GFP in the core of BFA compartments (Figure 1D, arrow heads). Then we tested the localization of BOR1-GFP to endomembrane compartments with secretory and recycling functions (Figure 1E–1H). BOR1-GFP-positive punctate structures showed close proximity to both endomembrane compartments labeled with mCherry-RabE1D and mCherry-RabA5D under control conditions (Figure 1E and 1G), whereas BFA treatment significantly promoted the colocalization between BOR1-GFP and these markers (Figure 1F and 1H). Additionally, some BOR1-GFP-positive punctate structures showed colocalization with mCherry-RHA1, an MVB/LE marker, under control conditions (Figure 1I). After BFA treatment, the mCherry-RHA1-positive MVB/LEs showed punctate structures in the periphery of BFA compartments (Figure 1J). To quantify the colocalization, we calculated the Pearson’s correlation between GFP and mCherry/mRFP fluorescence (Figure 1K). Pearson’s correlation values of BOR1-GFP and mCherry-Got1p homolog were 0.26±0.09 and −0.05±0.09 in control and BFA-treated root cells, respectively. Thus, BFA treatment clearly separated the Golgi apparatus and Golgi-associated organelles in which BOR1-GFP was preferentially localized. Pearson’s correlation between BOR1-GFP and VHA-a1-mRFP showed the highest value (0.77±0.08) among the markers in the control cells and was similar after the BFA treatment (0.82±0.04). Pearson’s correlations with mCherry-RabE1D were 0.41±0.04 and 0.84±0.04, in the control and BFA-treated cells, respectively. Those with RabA5d were 0.57±0.05 and 0.95±0.09, and with RHA1 were 0.30±0.14 and 0.58±0.08, in the control and BFA-treated cells, respectively. In BFA-treated cells, the highest correlation was observed with mCherry-RabA5D. Taken together, most BOR1-GFP proteins in the cytoplasm are localized in the TGN/EE and partially in other post-Golgi endosomes with secretory and recycling functions. These results support the view that BOR1 is constantly transported from the plasma membrane to early endosomes by constitutive endocytosis, and that it is recycled to the plasma membrane under low-B conditions.

**Figure 1.**
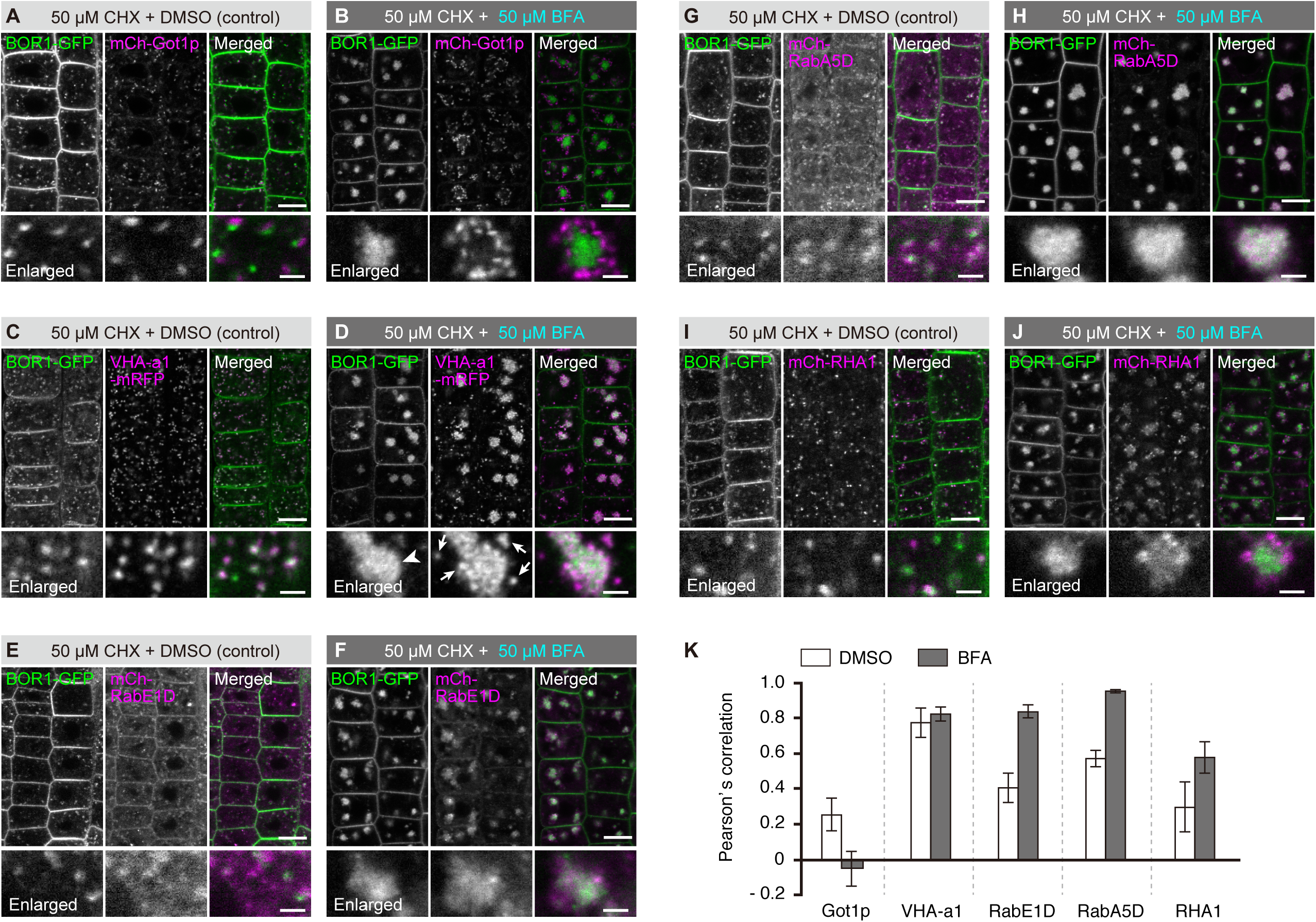
BOR1 is localized in the *trans*-Golgi network and transported via RabE1D and RabA5D positive endomembrane compartments under low-B conditions. (A–J) Confocal images of BOR1-GFP and endomembrane markers, mCherry-Got1p homolog (A and B), VHA-a1-mRFP (C and D), mCherry-RabE1D (E and F), mCherry-RabA5D (G and H), or mCherry-RHA1 (I and J) in the root epidermal cells of Arabidopsis seedlings grown on low-B medium (0.5 µM boric acid). Plants were incubated in low-B medium containing 50 µM CHX for 30 min followed by DMSO (A, C, E, G, and, I) or 50 µM BFA (B, D, F, H, and J) for 90 min. Lower panels represent enlarged images. Scale bars represent 10 (upper) and 2 µm (lower, enlarged images), respectively. (K) Pearson’s correlation between BOR1-GFP and endomembrane markers in the cytoplasm was calculated by the PSC colocalization plugin, Fiji/ImageJ. n = 10 cells. Error bars represent mean SD.

### 3.2. BOR1 is transported into the MVB/LEs labeled with the putative ESCRT-0 component TOL5

Generally, ubiquitinated membrane proteins undergo vacuolar sorting by the ESCRT machinery. TOL proteins are considered to have a redundant function as ESCRT-0 in the first step of sequestration of ubiquitinated cargoes in Arabidopsis (Gao et al. 2017). Among the TOL proteins, TOL5 was reported to localize to the cytosol, uncharacterized endomembrane compartments, and possibly the plasma membrane (Korbei et al. 2013). To examine whether TOL5 is involved in sorting of ubiquitinated BOR1, we performed colocalization analysis between BOR1 and TOL5. Although TOL5-mCherry showed strong cytosolic signals, partial colocalization with BOR1-GFP was observed in punctuate structures (Figure 2A). BFA treatment induced the separation of BFA compartments and TOL5-mCherry-positive punctate structures (Figure 2B). Pearson’s correlations between BOR1-GFP and TOL5-mCherry were 0.41±0.08 and 0.58±0.04 in the control (DMSO) and BFA-treated cells, respectively (Figure 2C). To further characterize the TOL5-mCherry-positive endomembrane compartments, plants were treated with 30 µM wortmannin (Wm), an inhibitor of the phosphatidylinositol 3-kinase (PI3K), that induces MVB/LE enlargement (Robinson et al. 2008), in addition to CHX and BFA (Figure 2D). Intriguingly, TOL5-mCherry showed ring-shaped structures that are independent of BFA compartments (Figure 2D), suggesting that TOL5 localizes to MVB/LEs. Furthermore, a high-B (100 µM boric acid) supply to plants treated with Wm triggered the internalization of BOR1-GFP into ring-shaped structure labeled with TOL5-mCherry (Figure 2E), indicating that BOR1 is transported into ILVs of TOL5-positive MVB/LEs under high-B conditions.

**Figure 2.**
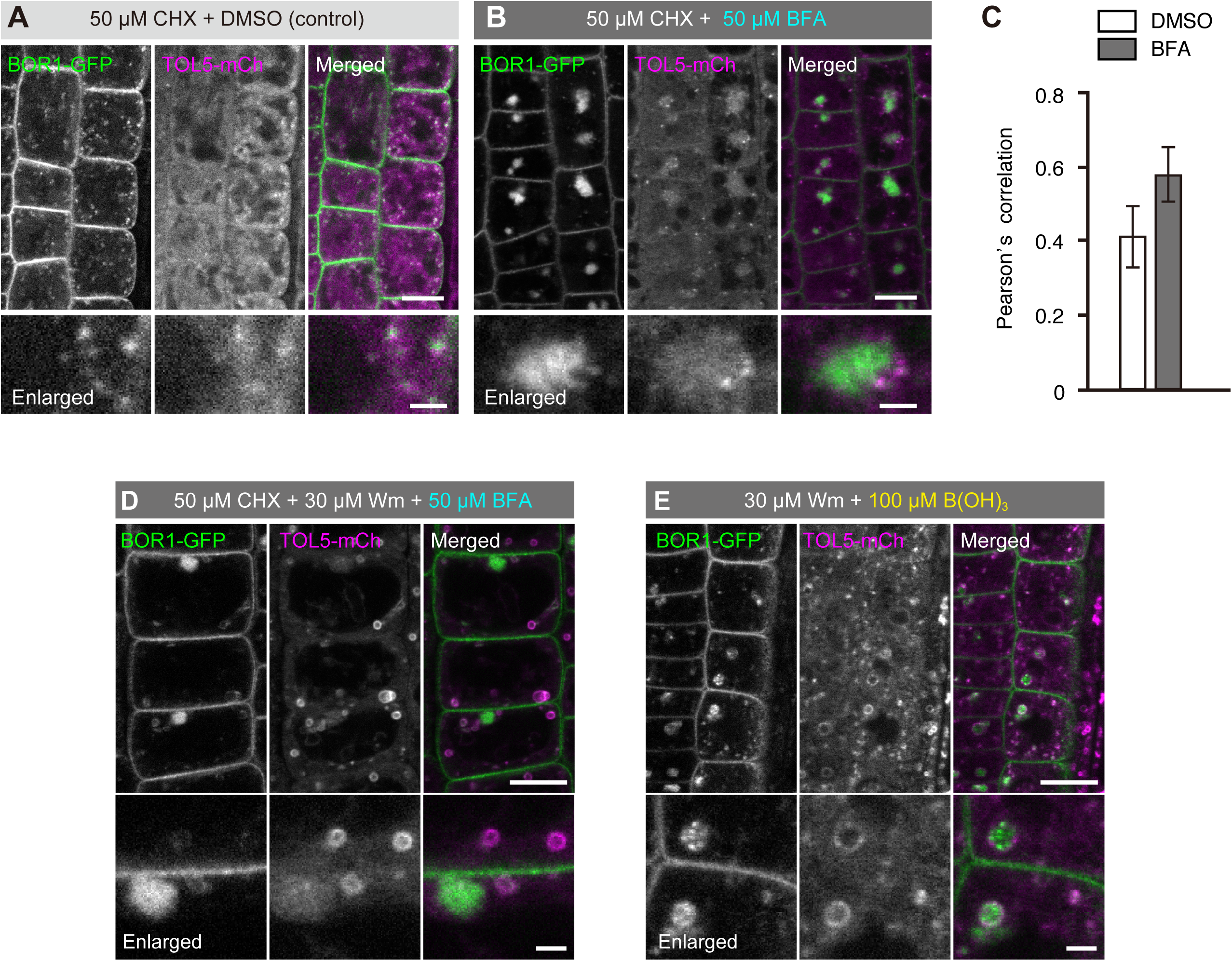
BOR1 is transported into TOL5-positive MVB/LE upon high-B supply. (A, B, D, and E) Confocal images of root epidermal cells of transgenic plants expressing BOR1-GFP and TOL5-mCherry under control of their own promoters grown under a low-B (0.5 µM boric acid) condition. Lower panels represent enlarged images. Scale bars represent 10 (upper) and 2 µm (lower, enlarged images). (A and B) The seedlings were incubated in low-B medium with 50 µM CHX for 30 min and then in medium with DMSO (A) or 50 µM BFA (B) for 90 min. (C) Pearson’s correlation between BOR1-GFP and endomembrane markers in the cytoplasm was calculated by the PSC colocalization plugin, Fiji/ImageJ. n = 10 cells. Error bars represent mean±SD. (D) The seedlings were incubated in low-B medium with 50 µM CHX for 30 min and then in the medium with 50 µM CHX, 30 µM Wm, and 50 µM BFA for 120 min. (E) The seedlings were incubated in low-B medium with 30 µM Wm for 20 min and then in medium with 100 µM boric acid and 30 µM Wm for 60 min.

### 3.3. TOL proteins are required for the vacuolar sorting of BOR1 but not for the polar localization and constitutive endocytosis of BOR1

We demonstrated that BOR1 is transported to the vacuole via TOL5-positive MVB/LE under high-B conditions. Here, to investigate whether functions of TOL proteins are involved in the vacuolar sorting of BOR1, we introduced the *pBOR1:BOR1-GFP* construct into a quintuple mutant of *TOL2, 3, 5, 6, and 9* genes, *tolQ* (Korbei et al. 2013). Although BOR1-GFP showed strong endosomal signals at 30 min in *tolQ* as well as wild-type plants, the BOR1-GFP signal in the plasma membrane remained for 120 min after high-B (100 µM boric acid) supply in the *tolQ* mutant (Figure 3). This result suggests that TOL proteins are required for efficient vacuolar sorting of BOR1 under high-B conditions.

**Figure 3.**
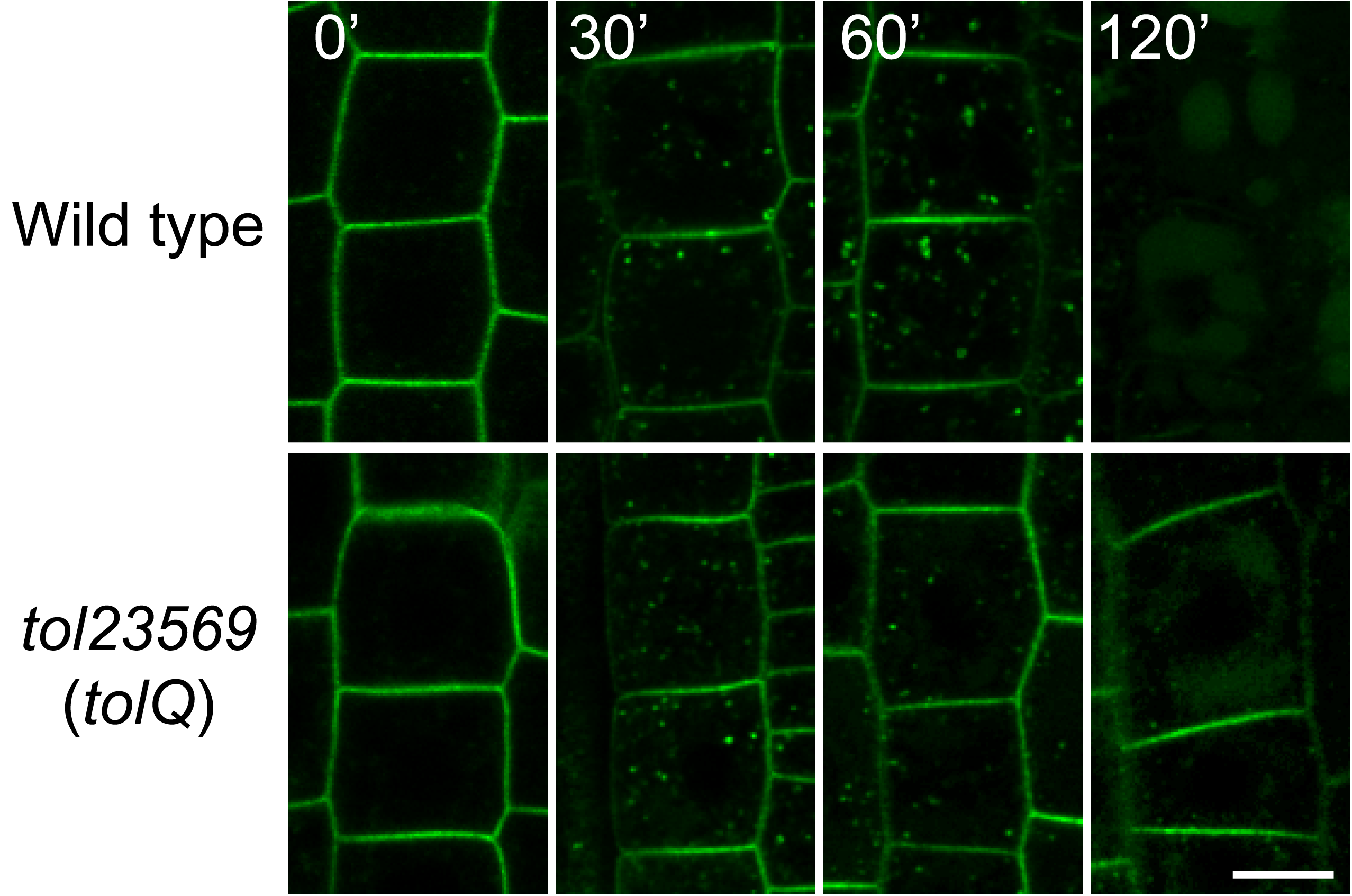
Vacuolar sorting of BOR1 requires TOL proteins-dependent MVB sorting. Confocal images of BOR1-GFP in the root epidermal cells of wild type (*bor1-1* background) and *tolQ* seedlings transferred from 0.5 to 500 µM boric acid conditions for the indicated time. Scale bars represent 10 µm. WT, wild-type.

In the *tolQ* mutant, BOR1-GFP was localized in the plasma membrane in a polar manner toward stele (Figure 4A), suggesting that TOL2, 3, 5, 6 and 9 are not involved in the maintenance of the polar localization of BOR1. Furthermore, in contrast to the ectopic localization of PIN2 in punctate structures (Korbei et al. 2013), the localization of BOR1 was not largely affected in the *tolQ* mutant (Figure 4A). To quantify the polarity of BOR1-GFP localization, we measured fluorescence intensities of BOR1-GFP at the plasma membrane in the wild type (*bor1-1* background) and the *tolQ* mutant and calculated the polarity index (Wakuta et al. 2015). Polarity indexes were 1.98±0.36 and 1.97±0.39 in the wild type and *tolQ*, respectively, with no significant difference (Figure 4B).

**Figure 4.**
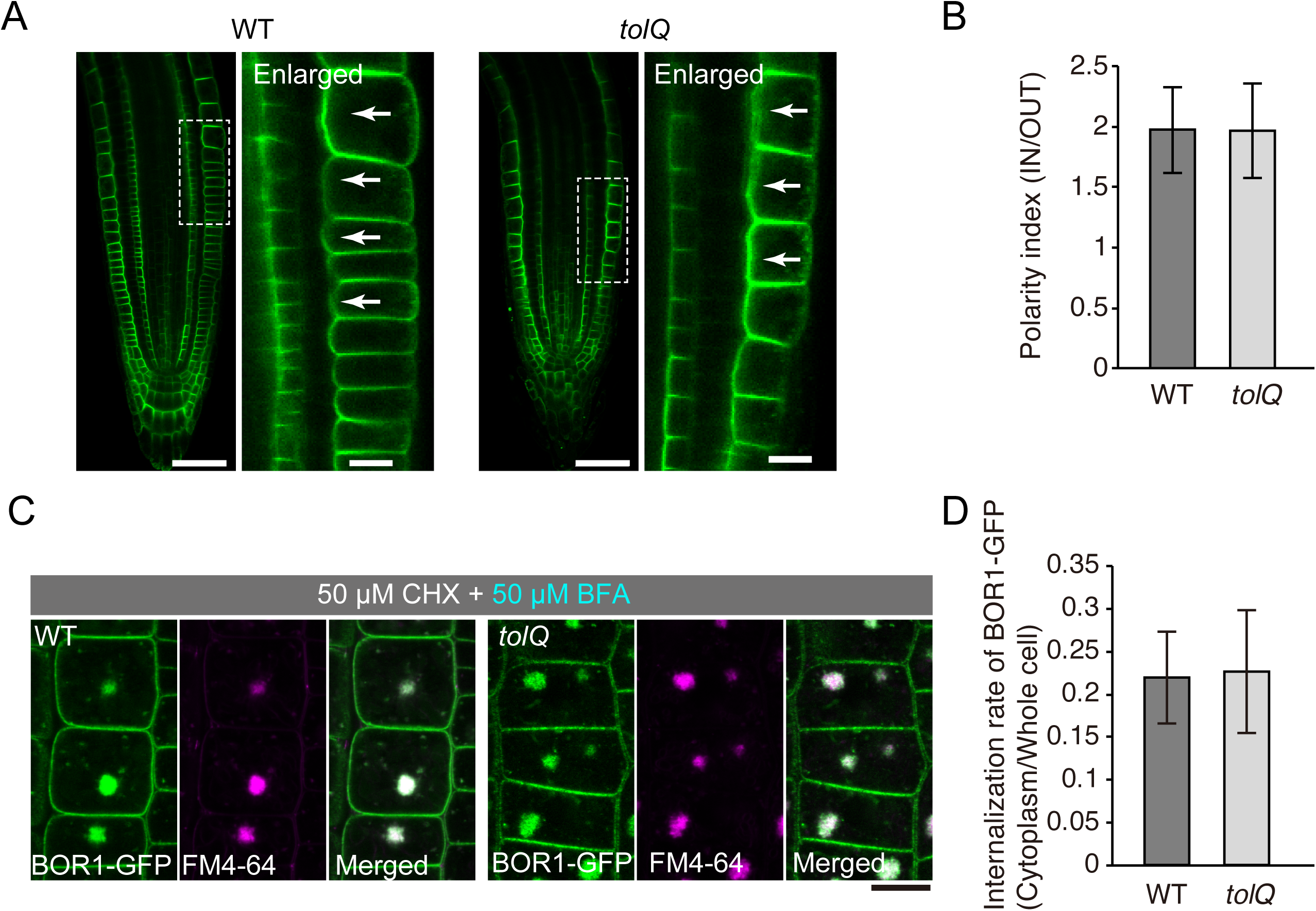
TOL proteins are not or little involved in the polar localization and the constitutive endocytosis of BOR1. (A) Confocal microscopy of BOR1-GFP in the primary root tips of wild type (*bor1-1* background) and *tolQ* seedlings grown on low-B medium. Right panels represent enlarged images of regions indicated in the left panels. (B) Polarity indexes of BOR1-GFP in wild-type and *tolQ* plants. Fluorescence signals in the inner-half of the apical and basal plasma membrane domains were divided by that in the outer-half. n = 40 (wild type) and 28 (*tolQ*) cells from three independent plants. (C) Confocal microscopy of BOR1-GFP in the epidermal cells of wild-type and *tolQ* plants incubated in low-B medium with 50 µM CHX and 2 µM FM4-64 for 30 min and then with 50 µM BFA and 50 µM CHX for 90 min. (D) Internalization rate of BOR1-GFP in wild-type and *tolQ* plants. n = 98 (wild type) and 124 (*tolQ*) cells from three independent plants. There was no significant difference by Student’s t-test. Error bars represent SD. Scale bars represent 50 (A, left panels) and 10 µm (A, right panels and C). WT, wild type.

We then examined the involvement of TOL2, TOL3, TOL5, TOL6, and TOL9 in constitutive endocytosis of BOR1 by using CHX and BFA. BOR1-GFP accumulated similarly in the BFA compartments of the wild type and *tolQ* plants at 90 min after BFA supply (Figure 4C). To quantify the internalization rate of BOR1-GFP, we calculated the ratio of fluorescent signals of BOR1-GFP in the cytoplasm and plasma membrane (Figure 4D; Yoshinari et al. 2016). Internalization rates of BOR1-GFP in wild type and *tolQ* plants were 0.22±0.05 and 0.23±0.07, respectively, with no significant difference. Thus, the polar localization and constitutive endocytosis of BOR1 are regulated independently of TOL2, 3, 5, 6, and 9 functions.

## 4. DISCUSSION

BOR1 is polarly localized in the plasma membrane of various tissues under low-B conditions, and it is rapidly transported to MVB/LEs for vacuolar degradation in response to high-B supply (Yoshinari and Takano, 2017). In the present study, we investigated the distribution of BOR1-GFP in different endomembrane compartments under low-B conditions. BOR1-GFP showed the strongest colocalization with VHA-a1-mRFP, a TGN/EE marker, under low-B conditions. This observation is consistent with a previous immune-electron microscopic analysis (Viotti et al. 2010). The BFA treatment in the presence of CHX did not affect the colocalization rate between BOR1-GFP and VHA-a1-mRFP, while it induced a significant increased colocalization of BOR1-GFP with the secretory/recycling-associated endomembrane markers, mCherry-RabE1D and mCherry-RabA5D (Figure 1). A previous study demonstrated that dominant-negative or pharmacological inhibition of RabA GTPases cause mis-localization of BOR1-GFP to unknown endomembrane compartments (Choi et al. 2011; Li et al. 2017). These data suggest that BOR1 is constitutively endocytosed from the plasma membrane, transported to the TGN/EE, and recycled to the plasma membrane via RabE1D and RabA5D-positive endomembrane compartments under low-B conditions.

In the *tolQ* mutants lacking multiple TOL proteins the high-B induced vacuolar sorting of BOR1-GFP showed significant delay (Figure 3), whereas the constitutive endocytosis and the polar localization of BOR1-GFP under low-B conditions were not affected (Figure 4). These results support the view that TOL proteins function in sequestration of ubiquitinated cargo as ESCRT-0 proteins in plant cells (Korbei et al. 2013; Gao et al. 2017). The exact compartment in which ESCRT-dependent sorting takes place is controversial because subunits of ESCRT-I, which act at downstream of ESCRT-0, are reported to be localized in the Golgi apparatus, TGN/EE, or MVB/LE, in different studies (Gao et al. 2017). In this study, we showed that a subset of TOL5-mCherry is localized in the MVB/LE and partially colocalized with BOR1-GFP (Figure 2A, B, D). We also showed that after Wm and high-B supply, BOR1-GFP and TOL5-mCherry were localized in the ILVs and the limiting membrane, respectively, in the enlarged MVB/LEs (Figure 2E). These results imply that recognition and sequestration of ubiquitinated cargo such as BOR1 by TOL proteins take place at the limiting (outer) membrane of MVB/LE.

In summary, we demonstrated that BOR1 is localized in the TGN/EE and the secretory/recycling-associated endomembrane compartments in the cytoplasm under low-B conditions, and that it is transported to TOL5-positive MVB/LEs upon high-B supply. We also demonstrated that the ubiquitinated BOR1 proteins are targeted to the vacuole via the TOL-dependent ubiquitinated-cargo sorting machinery.

---

**Supplemental Figure 1.** BOR1 is localized to the endosomes stained by FM4-64. Scale bars represent 10 (top) and 2 µm (bottom).

## AUTHOR CONTRIBUTIONS

A.Y., B.K., and J.T. designed research. A.Y. performed experiments and analyzed data. A.Y. and J.T. wrote the article.

## ACKNOWLEDGEMENTS

We thank Tomoko Shimizu and Kayo Konishi (Hokkaido University) for skillful technical assistance and Marcel Pascal Beier and Takuya Hosokawa (Osaka Prefecture University) for critical reading of the manuscript. We are grateful to Karin Schumacher (Universität Heidelberg) for providing us with VHA-a1-mRFP seeds and Niko Geldner (Université de Lausanne) for providing us with Wave line seeds.

